# Revisiting the explicit-implicit additivity assumption in visuomotor adaptation

**DOI:** 10.1101/2025.10.20.683435

**Authors:** Yifei Chen, Jordan A. Taylor

## Abstract

Explicit aiming strategies have been shown to play an important role in visuomotor adaptation – enabling rapid improvements in performance and affording flexibility – but their interaction and downstream consequences on implicit recalibration processes remain hotly debated. While early work assumed these processes combined additively, recent studies have challenged this view. However, these studies may have overlooked subtle spatial and temporal dynamics, which could influence how explicit aiming and implicit recalibration interact. Recent research shows that implicit recalibration anchors to where a person aims their movements, with aiming strategies directly shaping their spatial development. Moreover, implicit recalibration operates across multiple timescales, with both temporally volatile and persistent components. To examine whether these factors mask the true relationship between explicit strategies and implicit recalibration, we conducted a visuomotor rotation task while carefully accounting for the interplay of spatial and temporal dynamics. After controlling for spatial dynamics (plan-based generalization) and temporal dynamics (forgetting), we found a strong relationship between explicit strategies and implicit recalibration. Despite finding this strong relationship, it appears to be sub-additive, which may result from simple methodological imprecision, the operation of additional but unobserved processes, or more complex nonlinear interactions between processes.

## Introduction

Sensorimotor calibration is essential for accurate and efficient motor execution of everyday activities. Traditionally, adaptation was thought to occur solely through implicit processes, operating unconsciously to adjust movement in response to changes of the body or environment <u>(Shadmehr et al. 1998; Corkin</u> <u>1968)</u>. More recently, explicit strategies – conscious, deliberate changes in motor planning – have been shown to play a larger role than previously thought, particularly in visuomotor rotation tasks <u>(Hegele and</u> <u>Heuer 2010; Benson et al. 2011)</u>. These tasks are ideal for studying the potential influence of explicit strategies because the perturbations and their verbalizable solutions are in the same coordinate space. One key demonstration of this came from the aim-report paradigm <u>(Taylor et al. 2014)</u>, where participants explicitly indicated their intended aiming location before movement execution. This paradigm provided a direct measure of an explicit strategy on a trial-by-trial basis, allowing implicit recalibration to be inferred via the subtraction of the participant’s reported aim and their executed reach (Figure 1a). It revealed that the stereotypical adaptation curve (e.g., power-law function) reflects the dynamic interplay between explicit strategies and implicit recalibration (Figure 1b). Because the initial aftereffects closely matched implicit recalibration inferred during training, albeit slightly smaller, it was taken as a validation of the aim-report method <u>(Taylor et al. 2014)</u>.

**Figure 1.**
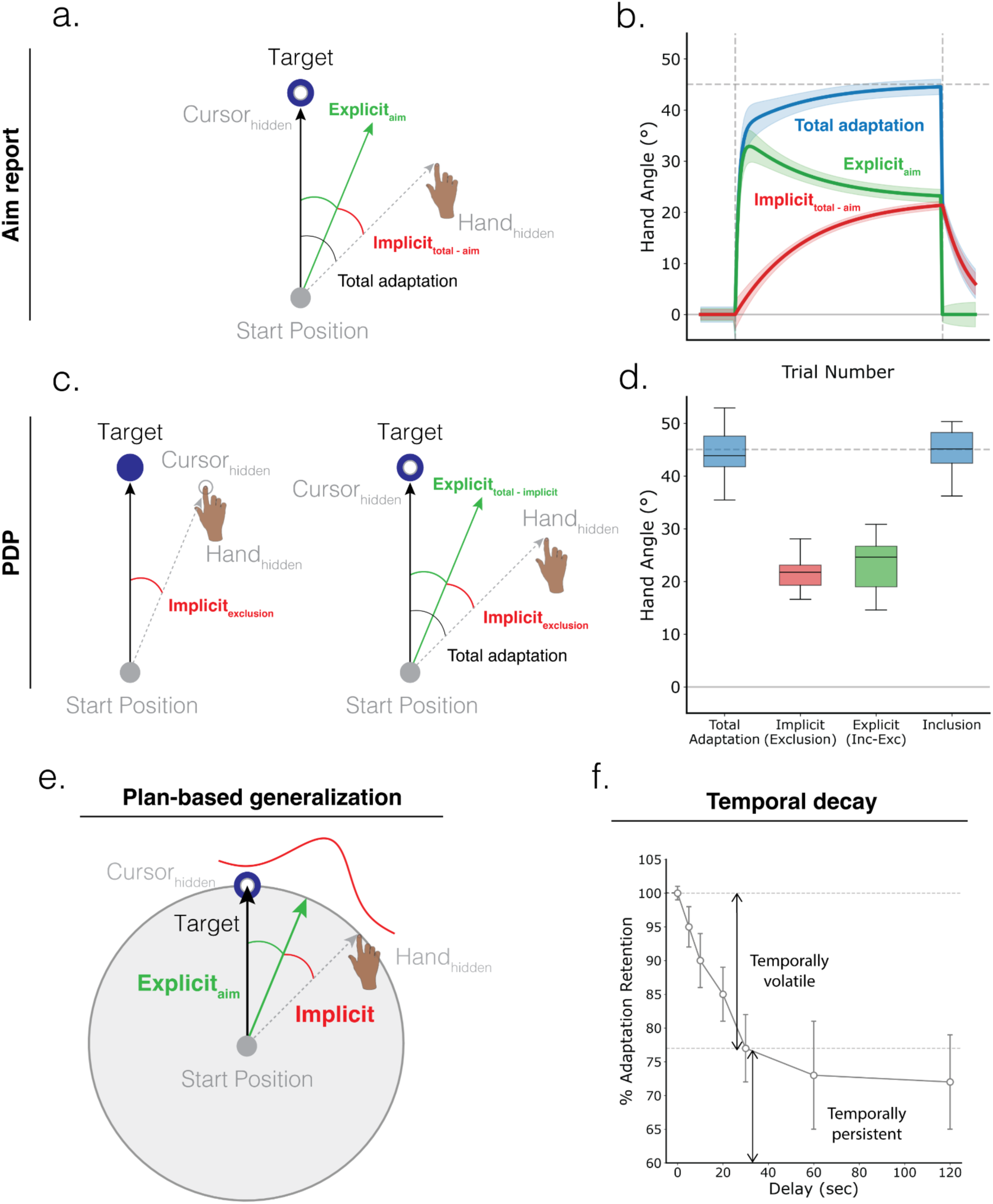
Different paradigms are used to dissociate implicit and explicit processes, and how plan-based generalization and temporal decay may affect the relationships between implicit recalibration and explicit strategy. Aim-report paradigm: (**a**) Explicit adaptation measured via movement intention reports; implicit adaptation calculated as total adaptation minus reported aim. (**b**) Provides trial-by-trial component measures. PDP paradigm: (**c**) Inclusion trials measure total adaptation while Exclusion trials isolate implicit adaptation through strategy suppression; (**d**) Explicit adaptation is derived from differences between Inclusion and Exclusion. (**e**) Plan-based generalization: Implicit adaptation peaks at the aiming location rather than the target location. (**f**) Temporal decay: Adaptation exhibits two components: a volatile component that nearly vanishes, and a persistent component that remains after 30 to 120s delay (recreated from Zhou et al. 2017).

The aim-report paradigm assumed linear additivity between these processes, which was weakly supported by correlations between aftereffects and implicit recalibration <u>(Taylor et al. 2014)</u>. Since then, many studies have tacitly adopted this assumption, though recent efforts to validate the method using the Process Dissociation Procedure (PDP) have raised questions about its accuracy <u>(Werner et al. 2015;</u> <u>Gastrock et al. 2020; Maresch et al. 2021; Modchalingam et al. 2019; Vachon et al. 2020)</u>. The PDP method was originally developed to distinguish between conscious and unconscious processes in memory and decision-making <u>(Jacoby 1991; Werner et al. 2015; Maresch et al. 2021; ‘t Hart et al. 2024)</u>. In the context of visuomotor adaptation, PDP dissociates explicit and implicit processes by instructing participants to suppress (Exclusion trials; Figure 1c, left) or apply any strategy (Inclusion trials; Figure 1c, right) they may have learned to counteract the perturbation. In theory, the hand angle observed at the final stage of training and Inclusion trials should match, and the difference between Inclusion and Exclusion trials should provide an estimate of the explicit strategy (Figure 1d). Despite these predictions, the measures of explicit and implicit do not appear to perfectly match, calling into question the methodological attempts to dissociate the different processes and, potentially, the assumption of linear additivity <u>(’t Hart et al. 2024)</u>.

Notably, both paradigms assume linear additivity, predicting that increases in implicit recalibration should result in proportional decreases in explicit strategy when total adaptation remains constant. Whether the relationship reflects indirect interaction between independent processes or competition between the processes remains an open question <u>(Albert et al. 2022)</u>. Evidence from aim-report and error-clamp studies suggests that implicit recalibration operates independently, proceeding in a stereotypical fashion regardless of error size or task relevance <u>(Bond and Taylor 2015; Morehead et al. 2017; Kim et al. 2018;</u> <u>Butcher and Taylor 2018)</u>. Under this view, explicit strategies simply compensate for the slack in implicit recalibration, achieving an angular value necessary for optimal performance. We should note that there is evidence for implicit recalibration also responding to changes in explicit strategies <u>(Miyamoto et al.</u> <u>2020)</u>, but these interactions could be explained by two relatively independent processes operating in series or a feedthrough arrangement <u>(Taylor and Ivry 2011)</u>. Alternatively, their inverse relationship could be the result of a more direct interaction where implicit and explicit processes compete for error information: if one consumes the error signal, less remains available for another <u>(Albert et al. 2022)</u>. In either case, an inverse relationship is assumed.

‘t Hart et al. (2024) tested the inverse relationship using both aim-report and PDP paradigms to obtain independent measures of each process to avoid mathematical dependencies inherent in each method <u>(Taylor et al. 2014; Albert et al. 2022)</u>. In the aim-report paradigm, implicit adaptation is derived by subtracting explicit aim reports from total adaptation, inherently creating a correlation between the two measures since one is mathematically dependent on the other. Likewise, in the PDP paradigm, explicit adaptation is inferred via subtraction of Exclusion from Inclusion. To test for a true correlation, at least two independent methods (e.g., aim-report and PDP) or, at least, data from two different phases of the experiment (e.g., PDP-Exclusion and washout phases) must be used. However, despite this dual-method approach, they found a lack of linear additivity using both independent and dependent measures of explicit and implicit processes, suggesting more complex interactions between the two. Interestingly, t’ Hart and colleagues failed to find a significant relationship even when one would be expected purely from mathematical dependency.

While these findings are concerning, potentially invalidating more than a decade of research, there have been several studies demonstrating subtle yet complex spatial and temporal interactions that could explain the lack of an observed relationship between implicit recalibration and explicit strategies <u>(Day et al. 2016;</u> <u>McDougle et al. 2017; Poh and Taylor 2019; Hadjiosif et al. 2023; Hadjiosif et al. 2024)</u>. First, plan-based generalization shows that implicit recalibration follows a Gaussian distribution centered at the explicit aiming location rather than the target location (Figure 1e; <u>Day et al. 2016; McDougle et al. 2017)</u>. During Exclusion trials, participants are instructed to reach directly to the target rather than their aiming location during training. Consequently, the measured implicit recalibration captures only the tail of the Gaussian distribution rather than its peak. Second, implicit recalibration has been shown to comprise temporally volatile and persistent components <u>(Hadjiosif et al. 2023)</u>, with the volatile component dropping significantly after a 30s to 1-min delay, leaving only the persistent component remaining (Figure 1f; <u>Zhou</u> <u>et al. 2017)</u>. The intertrial interval between successive reaches is often on the order of 5 seconds, and studies often have more than one target. Given typical intertrial intervals of a few seconds and multiple target locations, significant decay may occur between reaches to the same target. One or both of these factors, which arise from differences between training and PDP probe trials, could potentially explain the apparent lack of additivity between explicit strategies and implicit recalibration.

The present study tested the linear additivity assumption by independently measuring explicit and implicit processes while accounting for spatial (plan-based generalization) and temporal (decay) interactions. Using a within-subject PDP design, we manipulated target location and delay as participants adapted to a 45° visuomotor rotation, with one group reporting their aiming strategies and a control group without aim reports (to control for potential effects of the reporting procedure; see <u>Maresch et al. 2021; Hadjiosif and</u> <u>Krakauer 2021</u>). While the effects of plan-based generalization and temporal decay were observed only in the aiming group, we found a strong inverse relationship between explicit strategies and implicit recalibration across groups. However, the relationship was not perfectly additive, suggesting further methodological improvements are needed to accurately assess the relationship between the processes, additional but unobserved processes may be in operation, or more complex interactions between the processes.

### Materials and Methods Participants

Sixty participants were recruited from the research participation pool managed by the Department of Psychology at Princeton University in exchange for course credit. Two participants were excluded from analysis for not following instructions (see Data Analysis), resulting in a final sample of fifty-eight participants (37 females, 21 males; mean age: 19.64, SD: 1.24). Recruitment was limited to individuals who were right-handed, had normal or corrected-to-normal vision to receive visual feedback, and had normal color vision. Participants were assigned to either an aiming group (n = 29) or a control group (n = 29).

### Task and apparatus

All participants made rapid “shooting” movements with a digitizing stylus (Intuos 3, Wacom) to bring a virtual cursor (0.8 cm diameter) to a target (1 cm diameter) positioned 7 cm from the start position (Figure 2a). If the movement exceeded 300 ms, an auditory warning saying “too slow” was triggered. Visual stimuli were displayed on an LCD touchscreen monitor (Dell) mounted 23.5 cm above the tablet, preventing direct vision of the hand. Between trials, an empty circle dynamically adjusted in size based on the radial distance between the hand and the center of the tablet to guide participants back to the start position (1 cm diameter), where they held for 500 ms before proceeding. A desktop PC (Dell) running custom MATLAB software <u>(Brainard 1997)</u> controlled the task.

**Figure 2.**
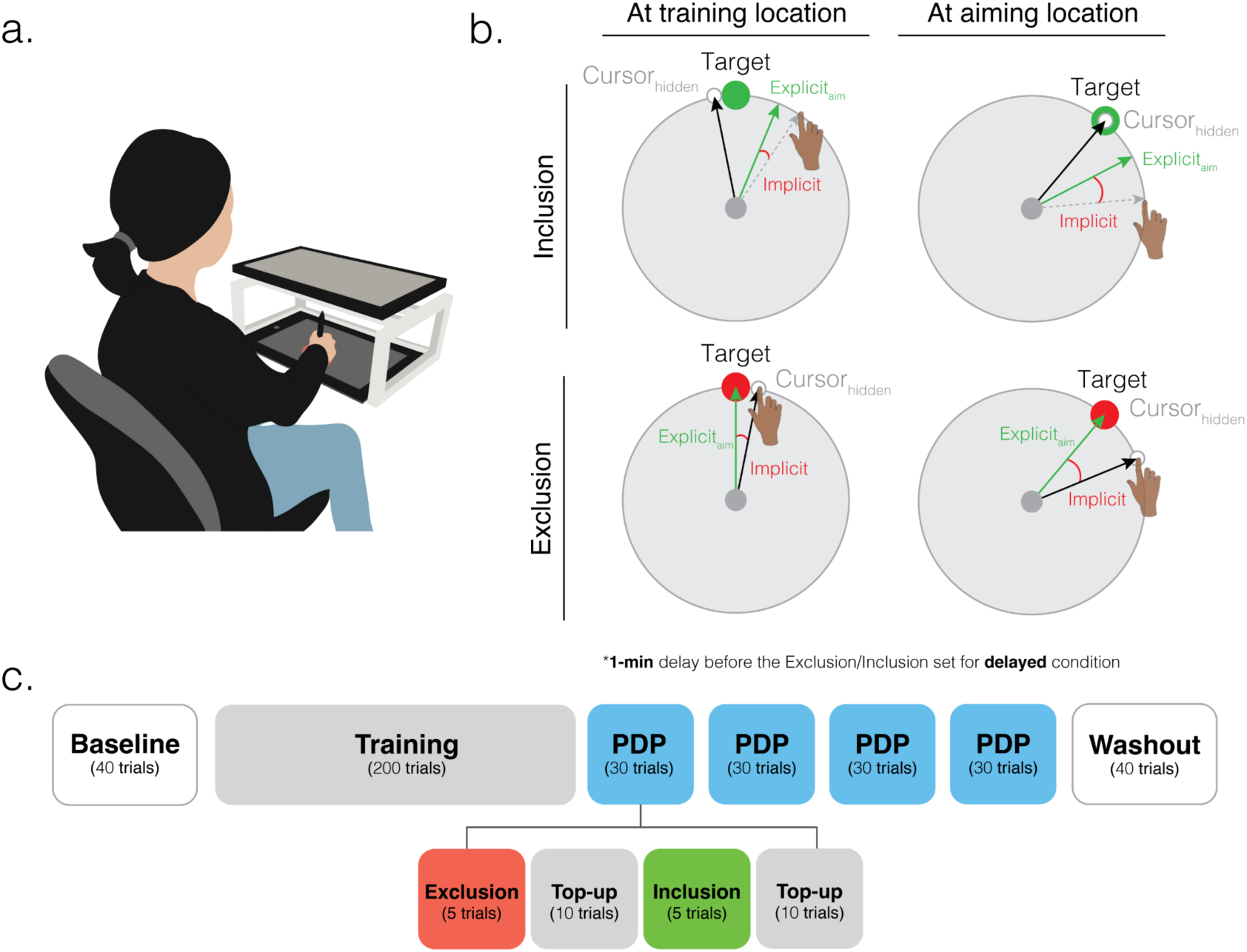
Experiment setup and design. (**a**) Participants performed fast-reaching movements by sliding a stylus on a digital tablet positioned under the monitor, which presented the visual workspace. (**b**) Example schematic of Inclusion trials (top) and Exclusion trials (bottom) for the target at the training location (left) and at the aiming location (right). (**c**) Experiment time course. The study employed a 2×2 within-subject design (Target Location: at training or at aiming locations × Delay: 1-min delay or no delay), resulting in four Process Dissociation Procedure (PDP) blocks.

The experiment consisted of 400 trials spread over four blocks. The first block was designed to familiarize participants with all aspects of the task, which consisted of 40 trials (Baseline block). For the first 10 trials, participants were provided with veridical continuous feedback of the cursor location. Cursor feedback was removed for the next 10 trials to measure any potential movement bias, before being reintroduced for an additional 10 trials with feedback. For the last 10 trials, the aiming group practiced reporting their intended aiming location with their left hand prior to executing each reach with their right hand, while the control group continued reaching without reporting their aims.

Following the baseline block, a 45° rotation was introduced between the movements of the hand and the visual feedback. Participants were then trained to overcome this rotation for 200 trials, while the aiming group continued to report their aim, and the control group did not (Training block). The direction of rotation was counterbalanced across participants. The location of the target was always at 90° (north) for the Baseline and Training blocks.

After the Training block, participants were introduced to the Process Dissociation Procedure (PDP), which unfolded over four phases of 30 trials (PDP blocks, Figure 2c). Each PDP block included 5 Exclusion trials, 10 Top-up trials, 5 Inclusion trials, and another set of Top-up trials. During Exclusion trials, cursor feedback was removed, and participants were instructed to reach directly toward the target without using any compensatory strategies (Figure 2b, bottom panels). The target turned from green to red to further indicate that participants should refrain from implementing any strategy. Exclusion trials are designed to assay implicit recalibration in relative isolation from explicit strategies <u>(Werner et al. 2015;</u> <u>Maresch et al. 2021; ‘t Hart et al. 2024)</u>. During Inclusion trials, cursor feedback was removed and participants were instructed to use any strategy they might have developed to counteract the perturbation in order to get their now unseen cursor on the target (Figure 2b, top panels). Inclusion trials are assumed to assay the joint operation of explicit strategies and implicit recalibration <u>(Werner et al. 2015; Maresch et</u> <u>al. 2021; ‘t Hart et al. 2024)</u>. In between the Exclusion and Inclusion trials were Top-up trials where cursor feedback was restored, and participants were instructed to try to get the rotated cursor on the 90° training target. These trials were designed to recover any adaptation that may have decayed during Exclusion/Inclusion trials since cursor feedback was removed. The order of the Exclusion and Inclusion sets alternated after each PDP block, and they were always separated by Top-up trials. Throughout the PDP blocks, the aiming group continued to report their aiming location by tapping on the monitor using their left hand.

The four PDP blocks (Figure 2c) were designed to test if plan-based generalization and temporal decay affect linear additivity. Here, we used a within-subject 2×2 factorial design with factors of Target Location (at training location or aiming location) and Delay (no delay or 1-min delay). In the Training Location condition, the target appeared at 90° (north), which is the same target location as during the training block, for both Inclusion (Figure 2b, top-left) and Exclusion trials (Figure 2b, bottom-left). In the Aiming Location condition, the target appeared at the average aiming angle from the five most recent Top-up/adaptation trials before the current Inclusion/Exclusion set for the aiming group (Inclusion example: Figure 2b, top-right; Exclusion example: Figure 2b, bottom-right). For the Control Group, the aiming target location was fixed at 39° based on the average late adaptation aim reports (last 10 trials of the Training block) from our pilot study <u>(Chen and Taylor 2025)</u>. To test for the effects of temporal decay, a 1-min delay was introduced between the final trial of Top-up and the onset of Exclusion/Inclusion in the Delay condition. The order of the four PDP blocks (Target Location and Delay combinations) was randomized for each participant. The experiment ended with 40 no-feedback no-aim-report trials to wash out any adaptation (Washout block).

### Data Analysis

Hand angles were calculated as the angular distances between the target and the hand’s endpoint position. Aim reports were calculated based on the angular distances between where participants tapped on the screen relative to the target. To standardize the data, we flipped all hand angles and aim reports in the same direction, ensuring that positive values always reflected the direction of counteracting the rotation, regardless of whether the rotation was clockwise (-) or counterclockwise (+).

To test the assumption of linear additivity, we examined whether implicit and explicit components exhibit an inverse relationship when total adaptation reached a similar level (45°). According to the strict linear additivity model, an increase in one component should correspond to a proportional decrease in the other, yielding a predictable slope. We modeled this relationship using linear regression:

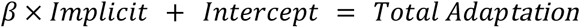

In the aiming group, where individuals reported their aims, explicit adaptation was quantified as the mean aim-reports from the five Inclusion trials, while implicit adaptation was calculated as the mean hand angle across the five Exclusion trials. In the control group, explicit adaptation was defined as the difference between the mean hand angles of the Inclusion and Exclusion trials, while implicit adaptation was assessed as the mean hand angle over the first five trials of the washout block. This approach ensured that explicit and implicit measures were based on non-overlapping, independent trial types in both groups.

To evaluate whether the observed slopes align with the linear additivity model, we first simulated data based on empirical data. Using the means and standard deviations of participants’ total adaptation and explicit aim reports during late adaptation (last 10 trials of the training block) from the aiming group, where we have a direct measure of explicit adaptation, we generated simulated data as follows:

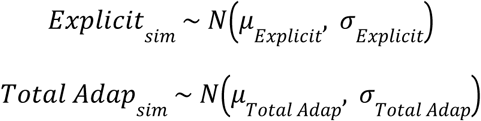

Next, we calculated simulated implicit adaptation using the linear additivity equation:

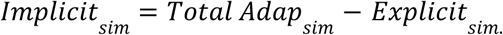

Finally, we fit a linear regression model to the simulated data to derive the predicted slope under the linear additivity model. This provided a benchmark for comparison with the observed slopes.

To examine how plan-based generalization (Target Location) and temporal decay influence implicit-explicit adaptation relationships and test the linear additivity model, we employed bootstrap hypothesis testing to compare regression slopes across experimental conditions. For each experimental condition, we estimated the slope relating implicit and explicit measures using ordinary least squares regression (OLS). Uncertainty was quantified with subject-level bootstrap resampling (1,000 iterations per condition): in each iteration, we resampled participants with replacement within each group and carried each selected participant’s data across all within-subject cells (Location and Delay) to preserve pairing for the aiming group. For every condition, we report the observed slope and the 95% confidence interval (CI) from the bootstrap distribution.

For the aiming group, we conducted a 2×2 factorial analysis examining how Target Location (Training vs. Aiming) and Delay (No-Delay vs. 1-minute) affected the slopes. For each bootstrap iteration, we computed marginal means and derived main effects as differences between factor levels and interactions as differences-of-differences across the appropriate cells. Statistical significance was determined by whether 95% bootstrap confidence intervals excluded zero, with two-tailed p-values calculated as twice the smaller tail probability. Significant interactions were decomposed using simple effects analyses. Note that the same analysis cannot be performed with the control group since the implicit measure remains the same for implicit adaptation across conditions (one washout block) while only the explicit measures (Inclusion -Exclusion) vary by condition. Comparing slopes across conditions would involve multiple comparisons using the same implicit data point, violating independence assumptions. Therefore, the slope for the control group is taken by averaging the implicit and explicit measures across all four conditions. Lastly, we compared overall slopes (collapsed across conditions) between groups and against simulated predictions from the linear additivity model using bootstrap confidence intervals.

As the dissociation and test of additivity between explicit and implicit critically hinges upon participants’ understanding of the specific instructions <u>(Chen and Taylor 2025)</u>, we first verified participants’ knowledge of the instructions. After being introduced to the task, they were asked to answer a few questions and would receive clarifying instructions if necessary before beginning the actual experiment. They were asked to indicate whether they should move directly toward the target or elsewhere in the following conditions: 1) baseline trials, 2) mismatch-on (mismatch between the cursor and the hand), and 3) mismatch-off trials. The correct answers were: 1) directly toward the target, 2) elsewhere, and 3) directly toward the target. Participants in the aiming group needed to answer an extra question about where to tap when reporting their aim (cursor vs. hand position), with the correct response being the hand position.

We then implemented a post-experiment survey to screen for individuals who failed to follow the instructions for both experiments. The post-experiment survey included 4 questions, each corresponding to a specific combination of Target Location (aiming location or training location) and trial type (Exclusion trial with a red target; Inclusion trial with a green target). The visuals matched those used in the actual task. For each question, participants indicated where they aimed by tapping along an “aiming ring” centered at the start position, with the target appearing on the ring. Note that the aiming location from the survey was the average aiming location from our pilot study <u>(Chen and Taylor 2025)</u>, which was the same aiming location for the control group during the actual task. Individuals who aimed within 0.8 cm of the target (the target’s diameter) during the Exclusion trials (both aiming and training locations) would pass the screening criteria and not be excluded, as aiming at a location other than the target in the Exclusion trials shows that they did not understand the instructions. Because Inclusion trials lack a dedicated instruction-check, we screened asymptotic performance by averaging the hand angle of each Inclusion block (the same trials used for the implicit–explicit correlation) across the four conditions and flagging participants whose mean hand angle was more than ±4 SD from the group mean. In the aiming group only, we applied the same ±4 SD criterion to the mean aim-reports to identify potential noncompliance or inconsistent reporting. As a result, two participants were excluded: (i) one control-group participant who failed to tap the target during an Exclusion trial when the target was at the aiming location, and (ii) one aiming-group participant whose aim-reports were extreme during an Inclusion block with the target at the aiming location.

All data and analysis code in Python can be openly accessed at: https://osf.io/zt8pd/overview?view_only=8b52f218257f4759838702974d040747.

## Results

### Task Performance

To investigate whether implicit and explicit adaptation linearly add up to total adaptation, we measured explicit and implicit processes while accounting for spatial (plan-based generalization) and temporal (decay) interactions. Specifically, the influence of spatial dynamics was examined by manipulating the Target Location (target at aiming vs training locations) and the influence of temporal dynamics by manipulating the Delay in cursor feedback (no delay vs 1-min delay) as participants adapted to a 45° visuomotor rotation in a within-subject design (n = 29). Importantly, to ensure that our assessment of the potential relationship between explicit and implicit processes was not the result of a necessary statistical correlation, we obtained independent measurements of each process via the aim-report and PDP techniques. We also included a control group (n = 29) who were not asked to report their aim on each trial to assess potential effects of the reporting procedure.

Participants successfully adapted to the 45° visumotor rotation, as indicated by an appropriate change in hand angle to counteract the perturbation over the last 10 trials of the Training block compared to Baseline in both aiming and control groups (Figure 3a; aiming: 43.75 ± 0.36°; *t*(28) = 65.73, *p* < 0.001; control: 43.15 ± 0.46; *t*(28) = 61.41, *p* < 0.001), and the adaptation is comparable across groups (*t*(56) = -0.61, *p* = 0.541). This comparable performance persisted throughout the PDP blocks, with no differences between groups in Exclusion (Figure 3b-c; *t*(56) = 0.69, *p* = 0.493) or Inclusion trials (Figure 3b-c; *t*(56) = 1.87, *p* = 0.067).

**Figure 3.**
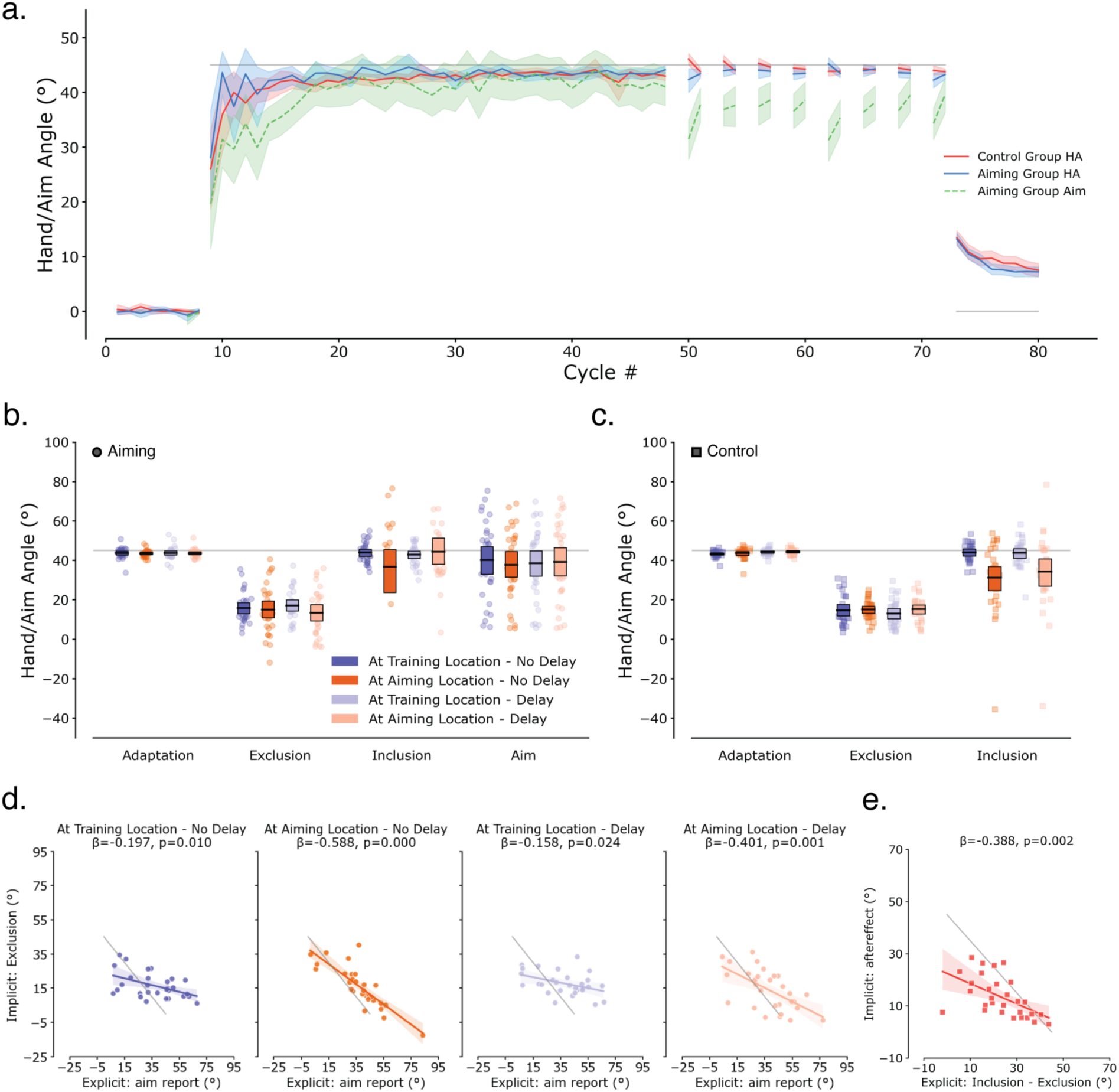
Significant but sub-additive relationships between explicit and implicit adaptation. (**a**) Time course of hand angle and aim report across trials. Solid lines show hand angle for aiming (blue, n = 29) and control (red, n = 29) groups; dashed green line shows aim report for the aiming group; grey line indicates the imposed rotation. Shaded regions represent 95% bootstrap CIs. (**b, c**) Hand angles during Top-up, Exclusion, and Inclusion trial blocks for (b) the aiming group with aim reports and (c) the control group without aim reports. Boxes indicate 95% CIs; dots show individual participants (circles: aiming group; squares: control group). (**d, e**) Relationship between explicit and implicit adaptation for (d) the aiming group across the 2×2 factorial design (Target Location × Delay) and (e) the control group collapsed across conditions. Colored lines show regression fits with the significance of the slopes (β) indicated. Grey diagonal lines represent the predicted inverse relationship under perfect linear additivity, where explicit + implicit = total adaptation.

Adaptation was maintained throughout testing, with no difference between Inclusion and Top-up trials when the probe-target was at the training location without delay (Figure 3b-c; aiming: Top-up, 43.82 ± 0.51°; Inclusion, 43.92 ± 0.99°; *t*(28) = -0.11, *p* = 0.912; control: Top-up, 43.34 ± 0.32°; Inclusion, 44.14 ± 0.87°; *t*(28) = -0.96, *p* = 0.35). Neither Target Location (aiming: *F* = 1.38, *p* = 0.249; control: *F* = 1.48, *p* = 0.234), Delay (aiming: *F* = 0.11, *p* = 0.748; control: *F* = 2.10, *p* = 0.158), nor their interaction (aiming: *F* = 3.51, *p* = 0.071; control: *F* = 2.41, *p* = 0.132) showed significant effects on Exclusion trials across groups. Similarly, Inclusion trials showed no effects of Target Location (Figure 3b; *F* = 0.68, *p* = 0.418) and Delay (*F* = 0.21, *p* = 0.647) in the aiming group. While there was no effect of Delay (Figure 3c; *F* = 1.21, *p* = 0.281) or an interaction (*F* = 1.82, *p* = 0.189), there was a significant effect of Target Location (*F* = 13.07, *p* = 0.001), with participants consistently reaching below the optimal hand angle when the probe target was at the Aiming Location (*t*(57) = -4.766, *p* < 0.001). While we did not find a significant effect of temporal decay (Hadjiosif et al. 2023; Zhou et al. 2017), it is important to note that the intertrial interval was relatively long on the first no-delay Exclusion/Inclusion trial (Supplementary Figure S1; 12.19 ± 5.25 s), which could have allowed significant forgetting on implicit adaptation to occur. Therefore, we suspect that the null finding here could reflect an absence of evidence rather than evidence of absence.

### Significant inverse relationships were found between implicit and explicit adaptation, with the effect of plan-based generalization and temporal decay observed in the aiming group

Although we found minimal impact of Target Location and Delay at the group level, our primary goal was to examine the relationship between implicit recalibration and explicit strategies at the individual level. To ensure independence between measures, we used non-overlapping trial types for each component. In the aiming group, explicit adaptation was quantified as the mean aim-report using the Inclusion trials, while implicit adaptation was the average hand angle across the Exclusion trials. In the control group, explicit adaptation was calculated as the difference between Inclusion and Exclusion trials, with implicit adaptation measured as the aftereffect.

In the aiming group, where participants explicitly reported their aiming directions, we found significant negative correlations between implicit and explicit adaptation across all conditions (Table 1; Figure 3d). This confirmed the expected inverse relationship — greater explicit strategy use was associated with reduced implicit recalibration. However, these correlations were consistently weaker than the slope of -1 predicted by linear additivity, suggesting that the two processes combine sub-additively rather than through simple summation. This pattern persisted even when analyzing only participants who strictly followed instructions (See supplementary Figure S2) using the same criteria from our pilot study, ruling out inattention or task misunderstanding as explanations <u>(Chen and Taylor 2025)</u>.

**Table 1.** Regression slopes between implicit and explicit adaptation across experimental conditions. Note: *β* represents the regression slope between implicit recalibration and explicit strategy. Under perfect linear additivity, the expected slope is -1.0. Bootstrap confidence intervals and *p*-values derived from 1,000 iterations.

| *Group* | *Target Location* | *Delay* | *Slope ($\beta$)* | *Confidence Interval* | *p-value* |
| --- | --- | --- | --- | --- | --- |
| *Aiming* | **Training** | **No** | -0.197 | [-0.352, -0.038] | 0.010 |
|  |  | **1-min** | -0.158 | [-0.270, -0.032] | 0.024 |
|  | **Aiming** | **No** | -0.588 | [-0.754, -0.482] | < 0.001 |
|  |  | **1-min** | -0.401 | [-0.604, -0.206] | 0.001 |
| *Control* |  |  | -0.388 | [-0.660, -0.146] | 0.002 |

Bootstrap hypothesis testing on the regression slopes revealed a significant main effect of Target Location, with stronger inverse relationships (more negative slopes) observed when the probe target was at the Aiming Location compared to the Training Location (Figure 5b; mean difference = -0.324, 95% CI [-0.458, -0.206], *p* < 0.001). And there is a marginal effect of Delay (mean difference = 0.116, 95% CI [0.000, 0.253], *p* = 0.052), meaning that when there was a delay, the relationship between the two components actually weakened slightly. However, there is no interaction between Target Location and Delay (mean difference = 0.152, 95% CI [-0.060, 0.368], *p* = 0.134). These results indicate that plan-based generalization and temporal decay, may significantly modulate the relationship between implicit and explicit adaptation when explicit strategies are independently measured through aim reporting.

The control group used the Process Dissociation Procedure without reporting aims, keeping the same 2×2 factorial structure as the aiming group. The parallel design allows us to isolate the effect of reporting aims and examine whether the same experimental manipulations would yield similar patterns when explicit strategies were measured through behavior rather than verbal report. Note, however, the design of the control group does not permit an independent measure of explicit aiming – there are no aim-reports – and the PDP method cannot estimate explicitly without using implicit from Exclusion trials, which creates a statistical dependency. As a result, we cannot perform the same factorial analysis as in the aiming group. To obtain a more generic estimate of the relationship for the control group, we quantified explicit adaptation by averaging the difference between Inclusion and Exclusion trials across all PDP blocks and quantified implicit adaptation from the single washout block at the end of training. When we averaged across conditions for the control group, we again observed a strong inverse relationship (Table 1; Figure 3e). The strength of the relationship (slope) did not differ significantly from the aiming group (Figure 4; mean difference = 0.027, 95% CI [-0.261, 0.313], *p* = 0.846).

**Figure 4.**
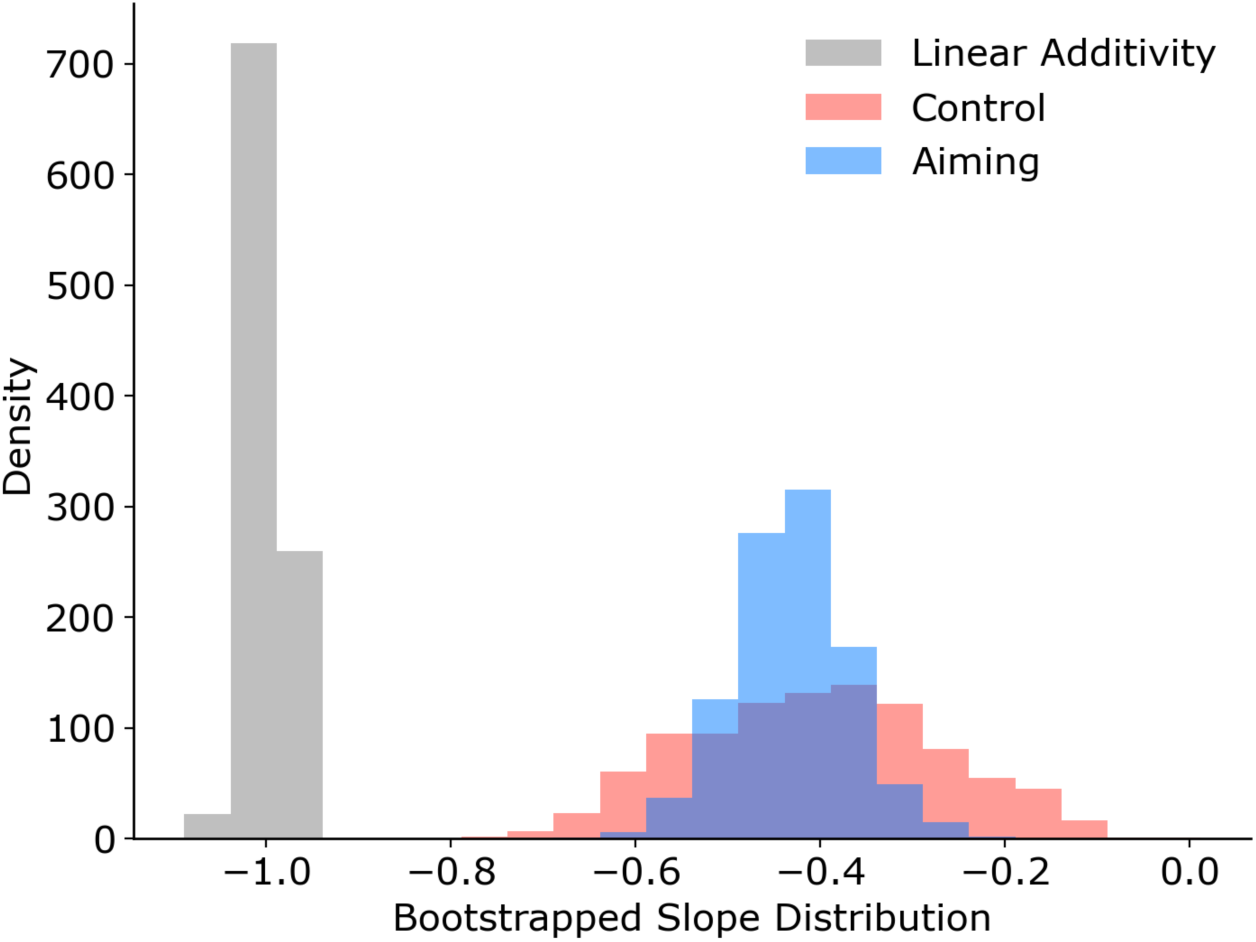
Distribution of bootstrapped slopes in aiming and control groups vs the linear additivity model. Colors denote different groups (gray = linear additivity; blue = aiming group; red = control group).

While using the difference between Inclusion and Exclusion trials has been the traditional way to compute explicit measures using the PDP paradigm, it may be problematic when targets appear at novel locations, as participants could become confused about how to express their learned adaptation during Inclusion trials. To test this possibility, we recomputed explicit adaptation using Top-up trials, where the target always appeared at the trained location, instead of Inclusion trials. This alternative approach yielded a much stronger inverse relationship (Supplementary Figure S3b; *β* = -1.061, *p* < 0.001, 95% CI [-1.303, -0.808]), approaching perfect linear additivity. This suggests that the traditional PDP approach may obscure the relationship between the two components when attempting to account for plan-based generalization by probing the adaptation at a new location.

To enable direct comparison between the two groups, we reanalyzed the aiming group data using the same methodological approach as the control group (aftereffects as the implicit measure, difference between Inclusion/Top-up and Exclusion as the explicit measure, averaging across conditions). Under these identical methodological conditions, the aiming group still showed sub-additivity between the two processes (Supplementary Figure S3c-d), suggesting that the act of reporting itself may indirectly or directly affect the relationship itself. We should note, however, that these analyses are post hoc and may be subject to the problem of multiple comparisons.

Finally, to quantify how much the observed relationships deviate from linear additivity, we conducted parameter recovery analysis, simulating slopes under the assumption of a linear relationship between explicit and implicit adaptation. Using variance estimates from our data, we simulated the distribution of slopes that would be expected if total adaptation were a direct sum of explicit and implicit components, subject to measurement noise. Bootstrap confidence intervals revealed that the observed slopes in both aiming and control groups differed significantly from the simulated slopes under the linear additivity model (aiming: mean difference = 0.569, 95% CI [0.442, 0.698], *p* < 0.001; control: mean difference = 0.596, 95% CI [0.337, 0.855], *p* < 0.001). Even if we restrict the analysis to only the aiming group when the PDP probes were at the Aiming Location without any delay, we find that the bootstrapped distribution of slopes still falls short of -1 (Supplementary Figure S4). Taken together, we find that there is a strong relationship between explicit and implicit adaptation, but it is sub-additive, challenging the simple linear additivity model of motor adaptation.

## Discussion

### Summary

Motor adaptation is not driven by a single, unitary process but instead emerges from multiple processes. Most taxonomies divide these learning and memory processes at the phenomenological level, with explicit and implicit processes at the top branch, with a variety of different subprocesses going down the branches <u>(Squire 2004; but see Wang et al. 2025)</u>. Along the explicit branch, there are at least two distinct components that have been identified: algorithmic-based (e.g., mental rotation) and retrieval-based (e.g., stimulus-response caching) strategies <u>(Pellizzer and Georgopoulos 1993; McDougle and Taylor 2019;</u> <u>Velázquez-Vargas and Taylor 2024)</u>. Along the implicit branch, a number of different components have been identified, such as implicit recalibration, use-dependent learning, proprioceptive recalibration, and reinforcement learning <u>(Cressman and Henriques 2009; Ostry et al. 2010; Izawa and Shadmehr 2011;</u> <u>Huang et al. 2011; Kim et al. 2019; Tsay et al. 2022)</u>. Given the variety and complexity of these underlying processes raises a fundamental question: how do they combine to produce overall adaptation? Many studies have assumed linear additivity at the top branch of this explicit-implicit taxonomy such that total adaptation is equal to the sum of explicit and implicit components, often estimating implicit as the difference between total and explicit adaptation <u>(Bond and Taylor 2015; Day et al. 2016; Mcdougle et al.</u> <u>2015; McDougle et al. 2016; Morehead et al. 2015; Taylor et al. 2014)</u>. But recent evidence challenges this view by showing non-additivity between the processes <u>(’t Hart et al. 2024)</u>.

Our study aimed to further investigate whether implicit and explicit adaptation combine linearly to explain total adaptation in motor learning while accounting for plan-based generalization and temporal decay, factors shown to influence the degree of implicit recalibration in complex ways. In a within-subject 2×2 design (Target Location: training vs. aiming; Delay: no-delay vs. 1-minute), we obtained independent explicit and implicit estimates using the Process Dissociation Procedure (PDP) in the control group and additional aim reports in the aiming group. While both groups demonstrated sub-additivity between explicit and implicit components, the aiming group revealed a more complex pattern. When adaptation was measured at participants’ actual aiming locations rather than at the target without a delay, the inverse relationship strengthened significantly, suggesting that spatial considerations fundamentally shape how these processes interact. However, even with this stronger coupling, the relationship remained sub-additive, falling short of the perfect trade-off predicted by linear models.

First, this could be simply due to the added complexity of the aim-report method causing confusion in the participants and, thus, affecting clean process measurement. Both the aiming report and the PDP method heavily depend on participants’ understanding of the instructions <u>(Chen and Taylor 2025)</u>. Second, given the number of learning processes that have been identified in recent years, it is likely that the aiming report and PDP method do not fully capture the contributions of these other processes. Finally, asking participants to explicitly report their aiming location or providing instructions to use or stop using compensatory strategies could fundamentally change the interaction between the myriad of explicit and implicit processes in potentially nonlinear ways.

### Spatial and temporal dynamics interactions

Failing to account for the effects of spatial and temporal dynamics on the relationship between explicit and implicit adaptation appeared to be a likely candidate for sub- or non-additivity, as it may have led to misinterpretations of a number of phenomena, such as spontaneous recovery, savings, and interference <u>(McDougle et al. 2017; Morehead et al. 2015; Day et al. 2016; Mcdougle et al. 2015; Schween et al.</u> <u>2018)</u>. While early work in motor adaptation assumed that adaptation would generalize around the target location <u>(Pine et al. 1996; Pearson et al. 2010; Brayanov et al. 2012; Taylor et al. 2013; Krakauer et al.</u> <u>2000; Tanaka et al. 2012)</u>, studies of force field adaptation suggested that generalization might instead be centered on the actual movement trajectory <u>(Gonzalez Castro et al. 2011)</u>. More recent studies of visuomotor rotations suggest that generalization occurs at the level of the movement plan, specifically at the aiming location rather than the target or the actual movement location <u>(Day et al. 2016; McDougle et</u> <u>al. 2017; Poh and Taylor 2019)</u>. In studies with a significant contribution of explicit aiming, plan-based generalization would predict an apparent reduction in implicit adaptation if it were probed at the training target during washout blocks or with the PDP method. As a result, additivity tests that infer implicit as the difference between total and explicit adaptation could appear sub- or non-additive because the probe is spatially misaligned with where learning is expressed.

To address this potential effect of plan-based generalization, we directly measured participants’ explicit aiming strategies and tailored our probe target to each individual’s Aiming Location to get a more accurate assessment of the implicit process. Indeed, we found that a significantly stronger inverse relationship between implicit and explicit processes emerged when implicit adaptation was measured at the Aiming Location. However, this relationship fell short of full additivity, suggesting that plan-based generalization alone could not account for previous reports of sub-additivity.

Temporal decay is another possible factor that could complicate the relationship between explicit and implicit processes. While explicit strategies remain stable, implicit recalibration decays over time: a volatile component that nearly vanishes within 1 min and a persistent component that remains stable <u>(Hadjiosif et al. 2023; Zhou et al. 2017)</u>. In studies where multiple targets were presented, implicit adaptation could potentially decay since it would take some time to get back to the same target <u>(’t Hart et</u> <u>al. 2024)</u>. Although the overall effect of temporal decay on adaptation was not significant – potentially due to the long pre-first-trial ITI that permitted forgetting (Supplementary Figure S1) – our result suggests that temporal decay slightly weakens the implicit-explicit relationship.

Even after aligning where and when learning is probed, the relationship between explicit and implicit components appears more complex than a simple linear sum. The modest but persistent deviation from additivity suggests that spatial and temporal factors, while important, cannot fully explain the sub-additive relationship between explicit and implicit adaptation. The remaining sub-additivity likely reflects the complex nature of adaptation: (i) methodological limitations in how we measure these processes, (ii) the operation of multiple unaccounted-for processes, or (iii) genuine interactions between learning systems.

### Current Methodological Limitations in Process Measurement

Our current methodologies fail to cleanly isolate explicit and implicit processes. Both the aim-report and PDP methods heavily depend on participants’ understanding of the instructions <u>(Chen and Taylor 2025)</u>, and even subtle differences in how these methods are implemented can yield divergent estimates of explicit and implicit contributions <u>(Maresch et al. 2021; Hadjiosif and Krakauer 2021)</u>. Our findings, also highlight important methodological considerations that can obscure the apparent relationship. Specifically, the null result from t’Hart et al. (2024) may stem from unclear instructions and an improper suboptimal measurement approach. In a pilot study, we found that participants’ understanding of task instructions is critical for accurate dissociation of learning components <u>(Chen and Taylor 2025)</u>. Without clear task comprehension, participants may conflate explicit and implicit processes or fail to engage them appropriately.

The aim-report procedure requires participants to introspect about their internal state and translate this into a verbalizable response, but the very act of questioning can alter what is being measured. When asked to report their “strategy,” participants face inherent ambiguity. Consider a participant adapting to a 45° rotation. They may report aiming 35° opposite to the rotation when asked about their strategy, yet they may only voluntarily control 30°. The 35° represents their reportable explicit knowledge: what they believe their strategy to be or what they think they are doing <u>(Maresch et al. 2021)</u>. The 30° represents their controllable explicit knowledge: what they can actually implement or suppress voluntarily <u>(Maresch</u> <u>et al. 2021)</u>. This 5° discrepancy cannot be attributed to simple motor noise; it reflects a fundamental dissociation between what participants think they are doing and what they can actually control. Should they report what they believe they are doing (35°), what they can actually control (30°), their intended full compensation (45°), or what they think the experimenter expects? The critical issue is that participants themselves may not have conscious access to these distinctions, and the ambiguity confounds the relationship between implicit and explicit processes when individuals are asked to report their explicit strategies.

In contrast, the PDP method seeks to minimally interfere by asking participants to express or suppress their strategy. This binary test bypasses much of the ambiguity of reporting, since it probes what participants can suppress rather than what they can articulate. However, the method raises limitations when accounting for plan-based generalization, as probing adaptation at a new location introduces additional complexity and confusion (e.g., the participant has never trained at that new location)

Moreover, the PDP approach suffers from not being able to provide an independent measure of explicit adaptation, leading to measurement limitations that can obscure true relationships between processes. These measurement challenges call into question studies claiming to find strong relationships between explicit and implicit adaptation. Albert et al. (2022) reported substantial correlations and proposed a competition model, but critically, they inferred both explicit and implicit components from the same behavioral output without independent dissociation. This approach necessarily guarantees correlations: when both measures derive from the same reaching behavior and one is calculated as the residual of the other, mathematical dependency can create artifactual relationships. The problem is compounded when plan-based generalization is ignored. If implicit adaptation is measured at the wrong spatial location, the explicit component calculated by subtraction will be overestimated, inflating correlations further. Without truly independent measures and proper spatial alignment, it becomes nearly impossible to determine whether observed correlations reflect genuine interactions or methodological artifacts.

Alternative experimental approaches have attempted to isolate specific components of adaptation. The error clamp method fixes the visual feedback at a constant angle regardless of hand movement, isolating implicit adaptation by eliminating trial-by-trial error correction that might drive explicit strategies <u>(Morehead et al. 2017; Kim et al. 2018)</u>. Gradual versus abrupt perturbation schedules have been proposed to differentially engage implicit and explicit processes, with the assumption that gradual rotations minimize awareness and explicit strategies while abrupt rotations provoke strategic aiming <u>(Saijo and Gomi 2010; Fernandez-Ruiz et al. 2011; Kagerer et al. 1997)</u>. Delay manipulations, which introduce temporal gaps between movement and feedback, selectively disrupt implicit adaptation while leaving explicit strategies largely intact, suggesting these processes have different temporal constraints <u>(Brudner et al. 2016; Schween and Hegele 2017)</u>.

Each of these methods seeks to privilege one process over another, with the hope that isolated components could be measured independently and then summed to reconstruct total adaptation. However, this isolation strategy encounters a fundamental problem: while these methods may successfully isolate processes at the group level, they necessarily eliminate individual differences in how these processes combine within a single person. For instance, one may utilize a more explicit strategy than implicit recalibration compared to another. At present, there is no cleanmethod for measuring explicit and implicit adaptation that avoids either some degree of measurement interference or requires generous inferential leaps about how isolated components relate to their interaction in natural learning contexts. Alas, there appears to be something akin to an Observer Effect where we cannot measure one process without (potentially) affecting or, at least, obscuring the other process.

### Multiple Implicit and Explicit Processes

Given the number of learning processes that have been identified in recent years, it is likely that the aim-report and PDP methods do not fully capture contributions of these other processes. Beyond the broad implicit versus explicit split, multiple sub-processes likely contribute within each system. On the implicit side, use-dependent learning creates repetition-induced biases that pull movements toward recently executed directions, even when error is minimized <u>(Diedrichsen et al. 2010; Huang et al. 2011)</u>. Proprioceptive recalibration shifts perceived hand position and the mapping between visual and proprioceptive estimates <u>(Cressman and Henriques 2009; Ostry et al. 2010)</u>. In addition, recent work proposes a distinct implicit component, implicit aiming, that biases action selection rather than execution *per se* and shows properties different from cerebellar-dependent recalibration (e.g., distinct temporal stability and contextual modulation), suggesting separable computations within implicit learning <u>(Wang et</u> <u>al. 2025)</u>.

Similarly, the explicit “system” likely encompasses multiple distinct processes rather than a single unitary strategy. Algorithmic strategies involve deliberate mental rotation and geometric calculation of aiming angles, requiring working memory and executive control <u>(McDougle and Taylor 2019)</u>. In contrast, retrieval-based strategies involve caching and recalling successful stimulus-response associations from previous trials, more akin to declarative memory retrieval than online computation <u>(Velázquez-Vargas and</u> <u>Taylor 2024)</u>. These two forms of explicit adaptation may have different dynamics, different capacity limitations, and potentially different relationships with implicit processes. A participant might simultaneously engage in both algorithmic and retrieval strategies, with their relative contributions shifting across trials. They might also employ higher-order rules about when to use algorithmic versus retrieval approaches. The fact that we collapse all of these controlled, deliberate processes under a single “explicit” estimate necessarily creates measurement error and could contribute to apparent sub-additivity.

The multiple subprocesses outlined here highlight, once again, the methodological limits of isolating distinct mechanisms within adaptation. The sub-additivity we observe could reflect contributions from a mixture of processes and other components that were not isolated by our measures. Without independent assays for each subprocess, it remains unclear whether their contributions are additive or whether interactions among them produce differences from additivity.

### Higher-order interactions between learning systems

Beyond the complexity introduced by multiple sub-processes, these systems at a more global level can potentially engage in dynamic interactions that lead to sub-additivity between the two systems. Albert et al. (2022) proposed that when explicit and implicit systems are driven by a common error signal, they compete rather than cooperate. Their competition model suggests that as explicit strategies increase to reduce target error, they simultaneously suppress implicit adaptation, not through simple subtraction but through inhibition. When one system, particularly the explicit system, becomes more active in correcting the error, it reduces the error signal available to the other system, thereby influencing its learning trajectory and observed behavior.

This competitive dynamic parallels findings from other learning domains. <u>Collins and Frank (2012)</u> showed that working memory (WM) and reinforcement learning (RL) systems interact in complex ways during instrumental learning. When WM successfully maintains stimulus-response mappings and successfully guides action selection, it reduces the signal available for the RL system. Conversely, when WM capacity is exceeded, RL must compensate, but does so less efficiently. This creates a fundamental tension where WM and RL dynamically share responsibility for action selection. Their relative contributions are weighted based on their estimated reliability or value, with the more reliable system having a greater influence on behavior at any given time. Working memory gates what gets learned through reinforcement, while RL signals influence what gets maintained in working memory.

Similar system-level interactions emerge between the hippocampus and basal ganglia, traditionally viewed as supporting independent declarative and procedural memory systems. Shohamy and colleagues have demonstrated that these systems engage in both competitive and cooperative dynamics depending on task demands <u>(Shohamy and Adcock 2010; Foerde and Shohamy 2011)</u>. During probabilistic learning, the hippocampus can actually interfere with basal ganglia-dependent habit formation by imposing explicit rules that override incremental statistical learning. In contrast, strong engagement of procedural systems can suppress hippocampal encoding of episodic details. These interactions are mediated by shared neuromodulatory signals, particularly dopamine projections from the midbrain that simultaneously influence both systems but with opposite effects depending on the specific receptors and circuits involved.

If implicit and explicit systems compete for error signals <u>(Albert et al. 2022)</u>, share dynamic control based on their capacity <u>(Collins and Frank 2012)</u>, and receive conflicting neuromodulatory signals <u>(Shohamy</u> <u>and Adcock 2010)</u>, then their contributions cannot simply be summed. Instead, engaging one system necessarily changes how the other operates. People with stronger working memory might show weaker implicit adaptation because their cognitive control systems more effectively suppress automatic recalibration <u>(Anguera et al. 2010; Christou et al. 2016)</u>.

### Challenges with Current Taxonomic Frameworks

The persistent challenges in demonstrating additivity highlight a more fundamental issue: we are trying to measure and categorize learning processes using taxonomic frameworks that may not align with how these processes are actually organized. The explicit-implicit distinction has a long historical tradition, originating from studies of patient H.M. and other cases of amnesia that revealed dissociations between conscious recollection and preserved motor skills <u>(Corkin 1968; Squire 2004)</u>. However, this distinction was originally developed for memory systems, not necessarily for the processes underlying motor adaptation. As studies have revealed increasing complexity within motor adaptation, multiple implicit/explicit processes and potential interactions between them, alternative frameworks have been proposed to better capture the underlying adaptation mechanisms. Yes, there remains no consensus on which framework best characterizes the space of learning processes.

One proposed alternative focuses on controllability rather than explicit awareness or knowledge As Maresch and colleagues (2021) proposed that we should differentiate between reportable explicit knowledge (the ability to describe the strategy being applied) and controllable explicit knowledge (the ability to deliberately choose to implement it). These reflect different aspects of conscious motor control and should not be treated as interchangeable. Reconceptualizing the framework around controllable versus uncontrollable components offers certain advantages. It avoids some of the ambiguity between explicit and implicit by focusing on what participants can actually modulate in behavior rather than what they can verbalize. This distinction may align better with experimental manipulations like the PDP, which operationalizes implicit learning as what persists when participants try to suppress their learned behavior.

Another alternative framework organizes processes by their goals: action selection versus action execution <u>(Wang et al. 2025)</u>. This framework distinguishes between processes that determine which action to perform – where to move – in order to achieve a task goal (action selection) versus processes that ensure the selected movement is accurately implemented (action execution). Critically, action execution is hypothesized to be intrinsically implicit, operating in an automatic, obligatory manner (e.g., implicit recalibration, use-dependent learning at the execution level). In contrast, action selection can engage both explicit processes (e.g., deliberate strategic aiming) and implicit processes (e.g., implicit aiming, habits). This framework offers explanatory power for previously conflicting findings. Processes like implicit aiming that contribute to action selection show savings and contextual sensitivity, while processes like implicit recalibration that contribute to action execution show anti-savings and are context-insensitive. These distinct patterns reflect fundamentally different computational goals – flexible selection to optimize task performance versus automatic execution to maintain calibration – rather than simply different levels of explicit awareness. This proposed action selection and execution framework shares some conceptual overlap with the controllable–uncontrollable framework, but they place the distinction at different levels and viewpoints: the former emphasizes the goal of what is being learned (selection vs. execution), whereas the latter emphasizes how that learning is expressed or modulated.

At present, there is no consensus on the optimal taxonomic framework for characterizing motor adaptation. The explicit-implicit distinction remains the most commonly used framework, partly due to historical precedent and the availability of measurement techniques designed around this dichotomy. But anyframework will begin with different assumptions about what matters most, which in turn shapes the experimental manipulations employed and how the results are interpreted. . Future work will need to develop better conceptual frameworks and measurement approaches that can capture the multi-dimensionality and complexity of motor learning processes.

## Conclusion

While our findings are consistent with previous reports on sub-additivity between explicit and implicit adaptation, we suspect that this sub-additivity is likely due to the operation of other learning processes that interact in interesting yet pedestrian ways, which are currently obscured by available experimental techniques and subject variance. Future efforts will require novel and careful methodologies to uncover if the multiplicity of learning processes truly adds up.

