## Supplementary Information for "Revisiting the explicit-implicit additivity assumption in visuomotor adaptation"

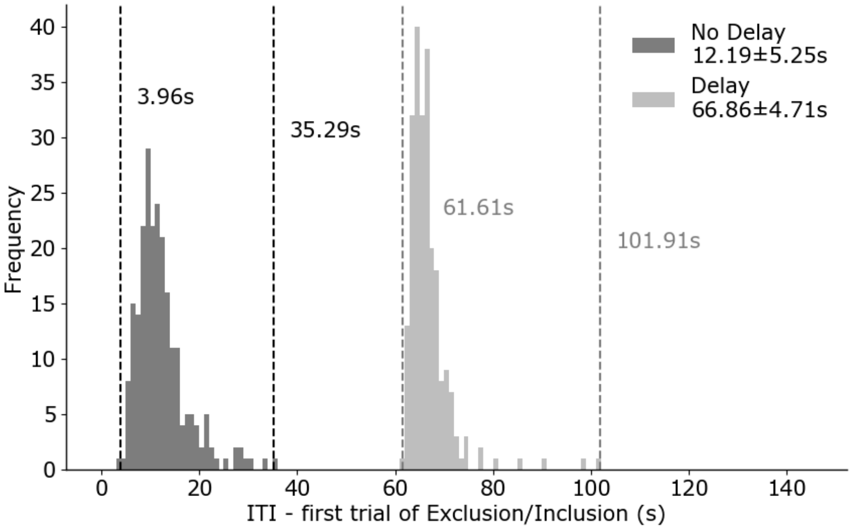


**Figure S1.** Mean intertrial intervals of the first Exclusion/Inclusion trial.

#### Sub-group analysis of the aiming group

To ensure that any lack of linear additivity was not due to instruction misunderstanding, we conducted subgroup analysis on the aiming group (n = 18) who demonstrated clear instruction comprehension based on three established criteria with slight modification (Chen and Taylor 2025). First, reported aims during Exclusion trials (instructed to aim directly at the target) had to fall within 0.8 cm of the target (the target's diameter), since aiming at a location other than the target in Exclusion trials shows that they did not understand the instructions. Second, aim reports from the last 5 trials of Top-up and Inclusion sets (without delay, at the training location) should not differ significantly, as these phases were identical except for the presence of cursor feedback. We should note that we did not consider Inclusion trials from PDP blocks where a delay was imposed or the target appeared at the aiming location, as differences in aim reports could stem from the change in conditions rather than from not following the instructions. Third, participants needed to demonstrate explicit strategy use, verified by a significant directional difference in aiming between the last 10 Training trials and Baseline trials. Notice that the subgroup analysis was not conducted with the control group since they did not report their aims throughout the task.

Similar to the full sample results, we found significant correlations when the target was presented at the aiming location (No Delay & Target at Training: $\beta = -0.166$, $p = 0.065$, 95% CI [-0.319, 0.009]; No Delay & Target at Aiming: $\beta = -0.562$, $p < 0.001$, 95% CI [-0.792, -0.412]; Delay & Target at Training: $\beta = -0.116$, $p = 0.156$, 95% CI [-0.252, 0.058]; Delay & Target at Aiming: $\beta = -0.300$, $p = 0.030$, 95% CI [-0.506, -0.035]). Nonetheless, the strength of the relationship is, again, below what would be expected from a purely linear combination (slope of -1), suggesting the lack of linear additivity within the aiming group is not due to misunderstandings of the instructions.

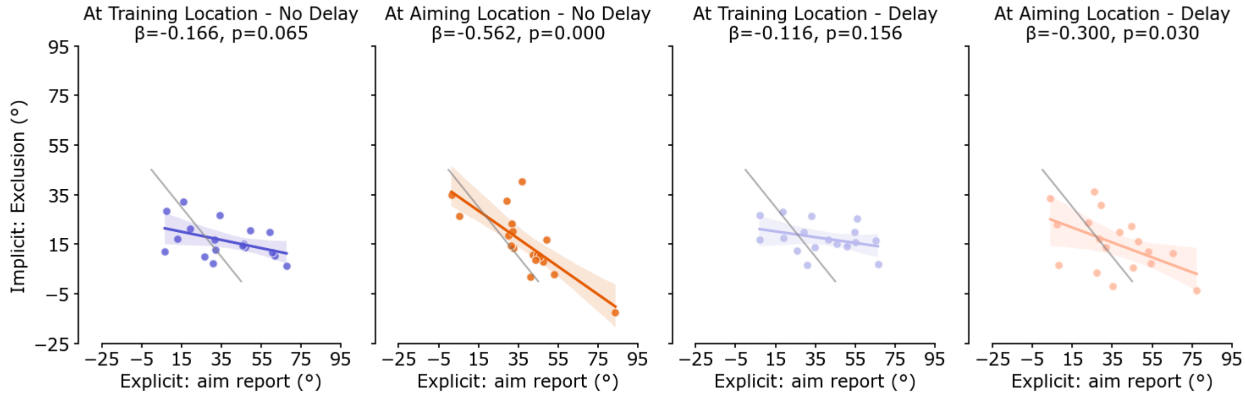


**Figure S2.** The relationships between explicit and implicit measures for the aiming subgroup that followed the instructions strictly (n = 18).

**Applying the same correlation analysis from the control group to the aiming group**

The two different approaches for control group style correlations analysis, using the differences between Inclusion/Top-up and Exclusion hand angles as a measure of explicit adaptation and aftereffect hand angle as a measure of implicit adaptation, were conducted as a direct comparison between the control and the aiming groups. Similar to what was observed in the original analysis, the aiming group shows significant inverse relationships but with shallower slopes than what would be expected based on the linear additivity model (Figure S3c; aftereffect vs. Inclusion - Exclusion: $\beta$ = -0.281, $p$ = 0.003, 95% CI [-0.418, -0.137]; Figure S3d; aftereffect vs. Top-up - Exclusion: $\beta$ = -0.482, $p$ < 0.001, 95% CI [-0.659, -0.318]).

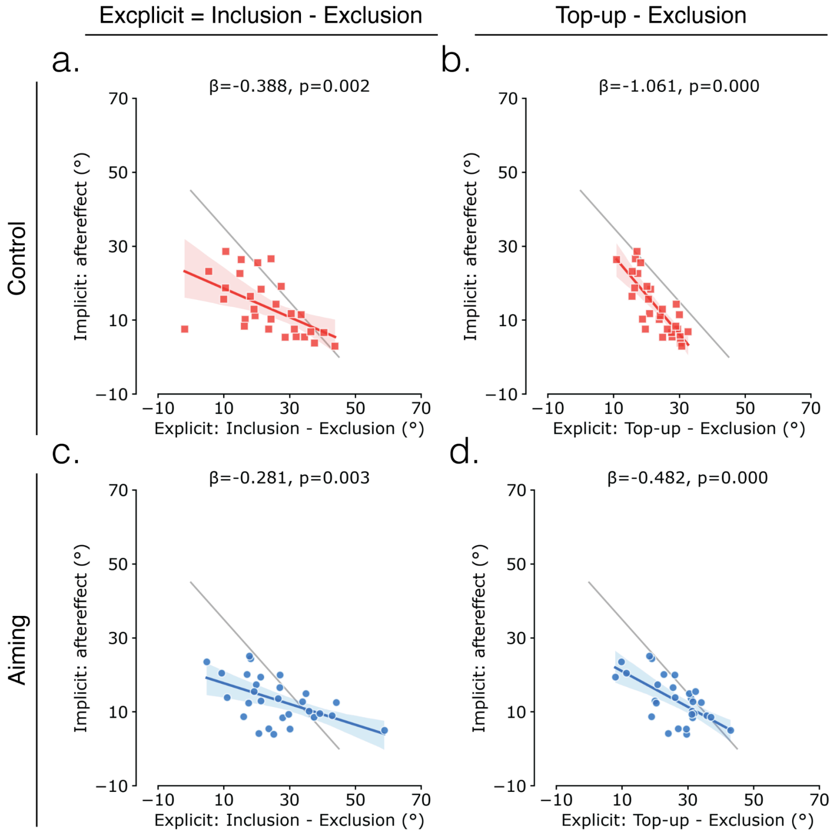

**Figure S3. Comparison of explicit-implicit adaptation relationships using different explicit learning measures.** Control group (top, n = 29) and aiming group (bottom, n = 29) data are shown for two explicit adaptation metrics: Inclusion minus Exclusion (left) and Top-up minus Exclusion (right).

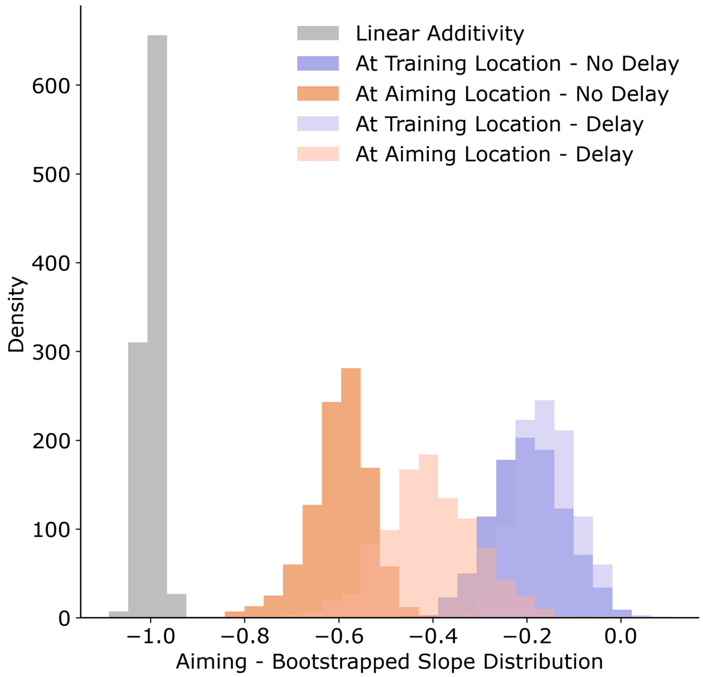


**Figure S4.** Distribution of bootstrapped slopes (aiming group, n - 29) across conditions vs the linear additivity model.

### Methodology Check

To validate the reliability of the Process Dissociation Procedure (PDP) and aim-report, we cross-checked whether different methods for assessing implicit recalibration, explicit strategy, and total adaptation yielded consistent results.

We first examined the accuracy of the PDP by comparing two key measures. First, we tested whether PDP Exclusion trials accurately captured implicit adaptation by comparing them to aftereffect measures at the training location with no delay. While PDP provided similar measures to aftereffect, the correlation fell short of perfect agreement (i.e., slope of 1) ($\beta = 0.725$, $p < 0.001$, 95% CI [0.610, 0.849]). However, a t-test shows that the measure of implicit adaptation is significantly higher using Exclusion trials compared to aftereffect trials ($t(57) = 4.429$, $p < 0.001$), likely due to the effect of plan-based generalization (while Exclusion trials measured implicit adaptation at both target and aiming location, aftereffect trials only measured implicit adaptation at the target location). We then compared total adaptation measures between Top-up trials and PDP Inclusion trials. Although no significant correlation was observed ($\beta = 0.091$, $p = 0.148$, 95% CI [-0.063, 0.267]), a t-test reveals no significant difference between the measures ($t(57) = 0.594$, $p = 0.555$), suggesting the lack of correlation may reflect the narrow range of hand angles rather than systematic differences.

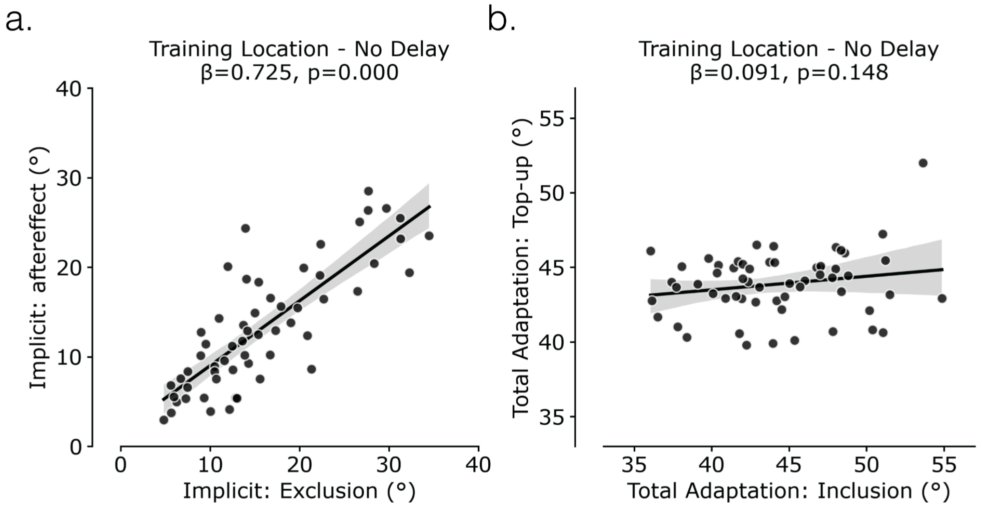


**Figure S5.** The relationships between (**a**) explicit measures using PDP Exclusion vs aftereffect trials and (**b**) implicit measures using PDP Inclusion trials vs Top-up trials across aiming and control groups (n = 59).

Within the aiming group, we examined whether explicit adaptation assessed through aim-report matches the one using PDP. We found significant and robust correlations across all conditions (No Delay & Target at Training: $\beta = 0.238$, $p = 0.005$, 95% CI [0.082, 0.386]; No Delay & Target at Aiming: $\beta = 0.953$, $p < 0.001$, 95% CI [0.635, 1.273]; Delay & Target at Training: $\beta = 0.254$, $p = 0.009$, 95% CI [0.078, 0.392]; Delay & Target at Aiming: $\beta = 0.584$, $p < 0.001$, 95% CI [0.241, 0.894]). The aiming location condition showed particularly strong agreement between the two explicit measures.

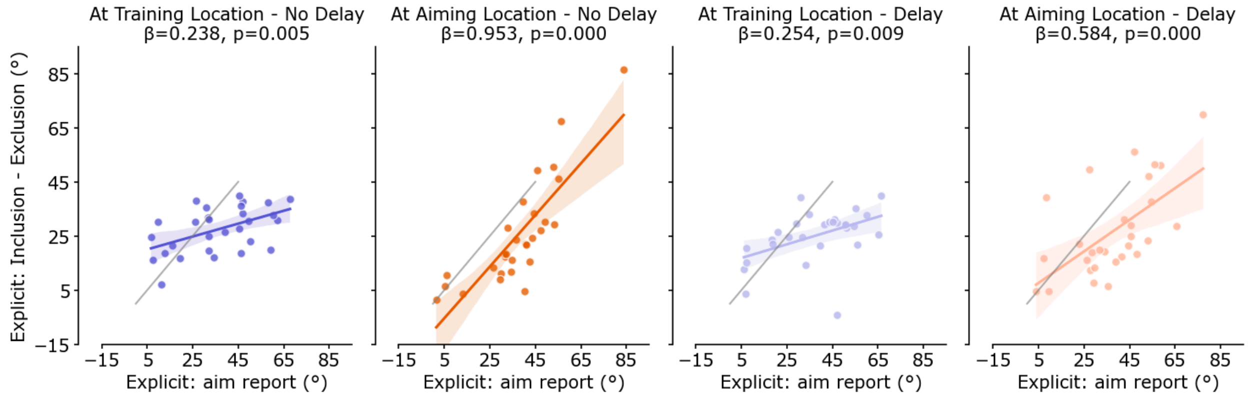


**Figure S6.** The relationships between explicit measures using aim-reports vs. PDP (difference between Inclusion and Exclusion) for the aiming group (n =29).

Overall, the analyses demonstrate reasonable consistency across measurement approaches, with PDP and aim-report providing relatively reliable estimates of both implicit and explicit components of motor adaptation, though some measures showed stronger convergent validity than others.
